# Multiplexed activation in mammalian cells using dFnCas12a-VPR

**DOI:** 10.1101/2021.01.31.429049

**Authors:** James W. Bryson, Jamie Y. Auxillos, Susan J. Rosser

**Affiliations:** Department of Quantitative Biology, Biochemistry and Biotechnology, University of Edinburgh, Edinburgh, United Kingdom; Centre for Synthetic and Systems Biology and UK Centre for Mammalian Synthetic Biology, School of Biological Sciences, University of Edinburgh, United Kingdom; Institute of Cell Biology, School of Biological Sciences, University of Edinburgh, Edinburgh, United Kingdom

## Abstract

The adoption of CRISPR systems for the generation of synthetic transcription factors has greatly simplified the process for upregulating endogenous gene expression, with a plethora of applications in cell biology, bioproduction and cell reprogramming. In particular the recently discovered Cas12a systems offer extended potential, as Cas12a is capable of processing its own crRNA array to provide multiple individual crRNAs for subsequent targeting from a single transcript. Here we show the application of dFnCas12a-VPR in mammalian cells, with FnCas12a possessing a shorter PAM sequence than As or Lb variants, enabling denser targeting of genomic loci. We observe that synergistic activation and multiplexing can be achieved using crRNA arrays but also show that crRNAs expressed towards the 5’ of 6-crRNA arrays show evidence of enhanced activity. This not only represents a more flexible tool for transcriptional modulation but further expands our understanding of the design capabilities and limitations when considering longer crRNA arrays for multiplexed targeting.

## Introduction

Synthetic transcription factors are modular proteins composed of DNA binding domains and transactivation domains, which enable up-regulation of targeted genes. Whilst a number of strategies have been developed (Becskei, 2020), the application of clustered regularly interspersed palindromic repeats (CRISPR) systems has greatly reduced the costs and complexity associated with generating synthetic transcription factors for targeting different loci (Pandelakis et al., 2020).

A hybridised CRISPR RNA (crRNA) and trans-activating crRNA (tracrRNA) enables targeting of the CRISPR associated protein 9 (Cas9) to a specific locus (Jinek et al., 2012). The spacer sequence within the crRNA confers target specificity, with binding and cleavage only occurring if the spacer sequence is complementary to the target sequence. There must also be a protospacer adjacent motif (PAM) sequence, which varies based on the Cas9 species of origin, adjacent to the target sequence. If the PAM sequence is present, then Cas9 can transiently melt the DNA to enable infiltration by the spacer sequence (Anders et al., 2014). Subsequently, if the spacer is complementary to the target sequence, then Cas9 will bind and cleave the target DNA.

The generation of a DNase inactive Cas9 variant (dCas9) has subsequently enabled the creation of RNA guided DNA binding domains, where the specificity of genome targeting can be altered by changing the 20 nt spacer sequence within a single guide RNA (sgRNA) composed of a fused crRNA and tracrRNA (Mali et al., 2013). A number of groups have subsequently generated synthetic transcription factors by fusing transactivation domains to dCas9 (Gilbert et al., 2013; Hilton et al., 2015). Cas9 derived synthetic transcription factors have been employed for a number of applications including; genetic circuits (Nakamura et al., 2019), cell reprogramming (Chakraborty et al., 2014) and biosensors (Krawczyk et al., 2020).

It is important to note that Cas9 represents only one of a variety of known CRISPR systems, with others possessing divergent and useful properties. In particular Cas12a/Cpf1, similarly to Cas9, functions as an RNA guided homing endonuclease. However, unlike Cas9, Cas12a can be targeted by a single crRNA (~40 nt) as opposed to requiring a combined crRNA and tracrRNA (~100 nt) (Zetsche et al., 2015). Furthermore, in contrast to Cas9, Cas12a possesses RNase activity and can recognise and process an array of adjacent crRNAs within a single transcript to enable targeting of the protein to multiple unique loci (Fonfara et al., 2016).

Whilst work has been carried out to generate synthetic transcription factors using DNase dead dCas12a variants from *Acidaminococcus sp*. BV3L6 and *Lachnospiraceae bacterium* ND206 (AsCas12a and LbCas12a respectively) (Tak et al., 2017), the related *Francisella novicida* variant (FnCas12a) has remained understudied, as initial work suggested it may not cut DNA *in vivo* (Zetsche et al., 2015). However subsequent work by Kim *et al*. showed it did possess activity in mammalian cells (Kim et al., 2016). FnCas12a was initially characterised as having a shorter PAM sequence than AsCas12a or LbCas12a *in vitro* (Zetsche et al., 2015). Subsequent cleavage assays in mammalian cells has shown a PAM sequence ‘K(G/T)Y(C/T)TV(A/C/G)’ enables optimal targeting for FnCas12a (Tu et al., 2017), compared to ‘TTTV’ for both AsCas12a and LbCas12a (Kim et al., 2017). ‘KYTV’ can on average be found every 21 nt. This targeting density is highly comparable to the targeting capacity of SpCas9 which has a PAM sequence ‘NGG’, which can on average be found every 16 nt. In contrast, the As/LbCas12a PAM sequence ‘TTTV’ can only be found on average every 85 nt.

The ability to target more synthetic transcription factors to a specific genomic region becomes essential in cases where narrow windows of targeting are optimal and in particular, when carrying out multiplexed targeting. One clear example is the case of gene network manipulation, where; 1) there is a 350 nt window within the promoter region where optimal transactivation is observed (Gilbert et al., 2014), 2) multiple promoters will be simultaneously targeted and 3) multiple studies including this work show that targeting more than one copy of the synthetic transcription factor to the same promoter can enable enhanced transactivation (Maeder et al., 2013; Tak et al., 2017).

In the following work we show that FnCas12a can be engineered and applied as a synthetic transcription factor in mammalian cells, before subsequently exploring whether dFnCas12a-VPR shows orthogonality when screened alongside dAsCas12a-VPR and dLbCas12a-VPR. We then test whether single crRNAs are sufficient for gene activation and look for synergistic transactivation when multiple crRNA target a single promoter. We further explore multiplexed activation from single crRNA arrays. Finally, we look into the role of position of targeting crRNA within 6-crRNA arrays on the capacity to transactivate targeted genes.

## Results

### 1 – dFnCas12a-VPR transactivates Luciferase plasmid reporter in mammalian cells

To assess the capability of different Cas12a systems to be adapted as synthetic transcription factors, three variants of Cas12a were chosen. The As, Lb and Fn variants were selected due to their well characterised nature (As and Lb) or the wide targeting range of the PAM sequence (Fn). DNase inactive variants were generated before the VPR transactivation domain was cloned onto the 3’end of each dCas12a.

The three variants (As, Fn and Lb) were initially screened alongside dCas9-VPR using a dual luciferase assay. Utilising a dual plasmid reporter system (Kleinjan et al., 2017), each of the dCas12a-VPR constructs and dCas9-VPR were targeted upstream of a Firefly luciferase gene using the respective crRNAs (dCas12a-VPR variants) or sgRNA (dCas9-VPR) (Figure 1A).

**Figure 1.**
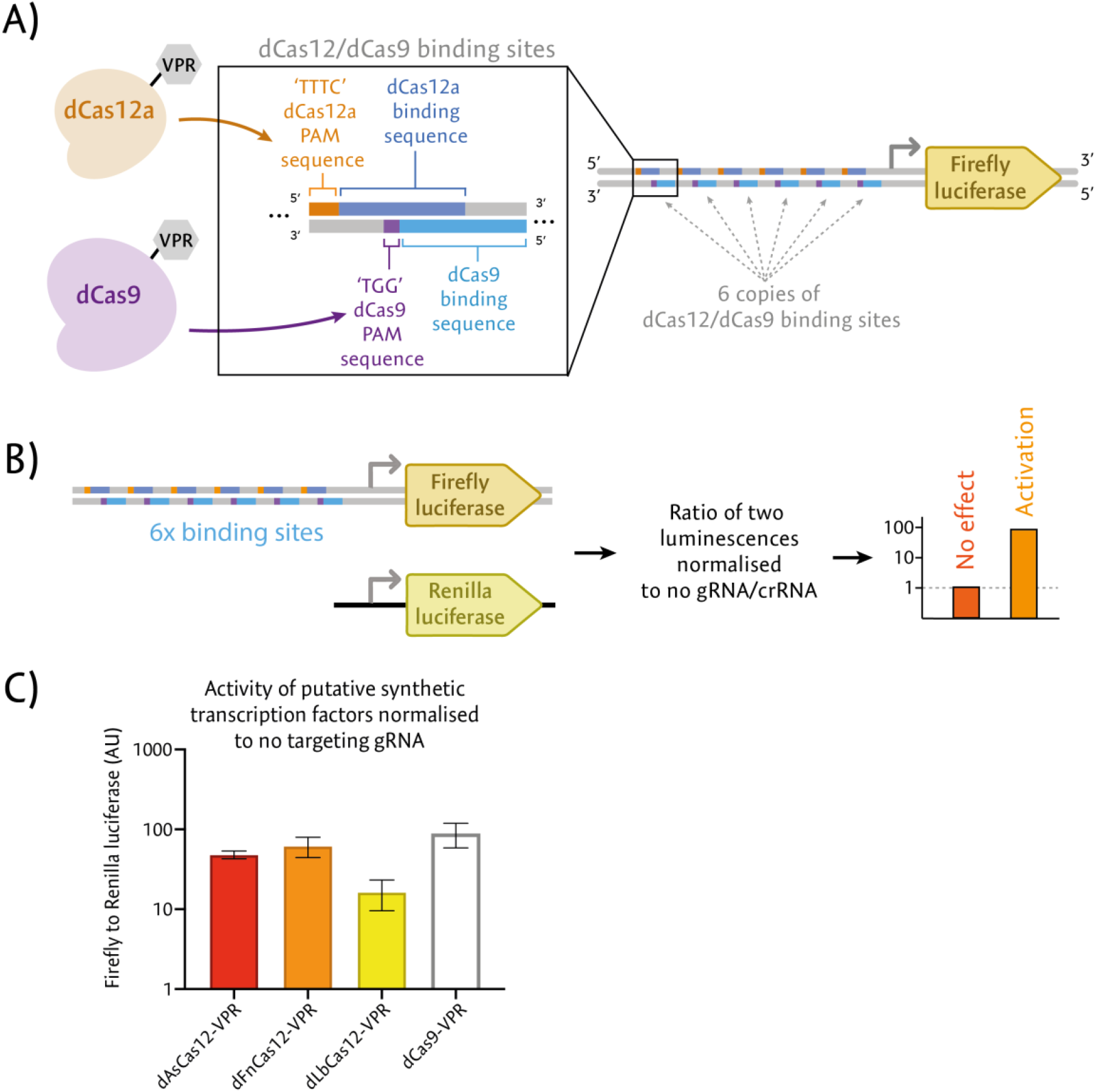
Screening dCas12a-VPR constructs using plasmid-based Firefly luciferase reporter. **A)** Schematic representation of the targeted promoter region within the Firefly Luciferase reporter plasmid. The top strand of the promoter region contains six repeated binding sequences for the dCas12a constructs (dark blue) with adjacent PAM sequences that can be recognised by all three variants ‘TTTC’ (orange). The bottom strand of the promoter region contains 6 repeated binding sequences for dCas9-VPR (light blue) with adjacent PAM sequences that can be recognised by dCas9-VPR (purple). **B)** Diagrammatic representation of the dual luciferase reporter assay. Alongside the targeted Firefly luciferase reporter plasmids, a non-targeted *Renilla* luciferase plasmid was delivered to enable normalisation of the relative Firefly luciferase activity between test and control conditions. If a putative synthetic transcription factor was able to transactivate the targeted Firefly luciferase gene, then the ratio of Firefly to *Renilla* luciferase activity would be increased compared to the negative control condition. **C)** Testing the three dCas12a-VPR variants alongside dCas9-VPR using the dual luciferase assay. Each construct is delivered with a targeting crRNA/gRNA and the resulting ratio is normalised to the ratio when delivered without a crRNA/gRNA. The results represent three biological replicates and the error bars display the SEM.

The plasmids expressing the synthetic transcription factor and crRNA/sgRNA were co-transfected alongside the targeted Firefly luciferase plasmid and a control *Renilla* luciferase plasmid (Figure 1B) into HEK293 cells. Two days post-transfection the ratio of the targeted Firefly luciferase to the normalising *Renilla* luciferase was measured for each variant. Of interest the Fn variant of dCas12a-VPR appeared to perform best when compared to dCas9-VPR (Figure 1C), showing significant transactivation (62 fold, P = 0.027 based on two-tailed students t-test).

### 2 – Orthogonality observed between dCas12a-VPR variants

We subsequently sought to test for orthogonality between the three dCas12a-VPR variants (Figure 2A). The activity of each variant when delivered with crRNAs from each of the variants or none was measured using the dual luciferase assay previously described (Figure 1B).

**Figure 2.**
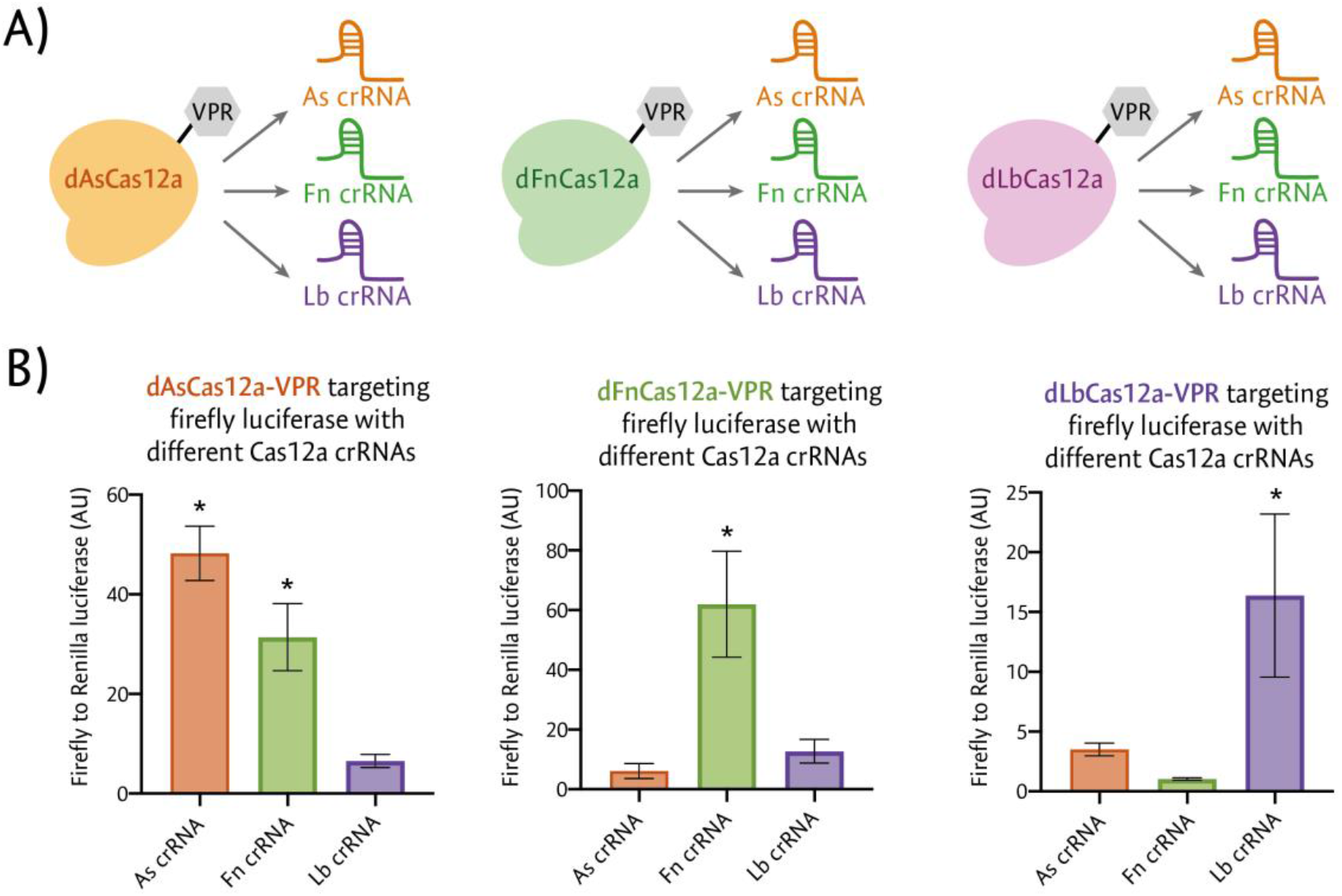
Testing for orthogonality between different dCas12a-VPR variants. **A)** Schematic representation of the testing of orthogonality using either native (e.g. dAsCas12a-VPR with As crRNA) or non-native (e.g. dAsCas12a-VPR with Fn crRNA) crRNA pairings. **B)** The three dCas12a-VPR variants (As, Fn and Lb) were screened for orthogonality using the dual luciferase assay. Each dCas12a-VPR construct was delivered with each of the three targeting crRNA (As, Fn or Lb) or no targeting crRNA. Results display the mean luciferase activity relative to no targeting crRNA from three biological replicates. Error bars show SEM and the stars (*) denote significant (P < 0.05) expression relative to no crRNA (based on a post hoc Dunnetts test).

We observe evidence of cross-reactivity between the As and Fn variants of dCas12a-VPR, with dAsCas12a-VPR targeted with the Fn crRNA showing significant transactivation (P = 0.003). Of interest the Fn and Lb variants do not show evidence of cross reactivity, suggesting they could serve as an orthogonal pair (Figure 2B).

### 3 – Single crRNAs are sufficient for transactivation of endogenous genes

Having demonstrated the activity of dFnCas12a-VPR using a plasmid-based reporter, we next sought to test whether transactivation of endogenous genes could be achieved. The three genes *HBB, ASCL1* and *IL1RN* were chosen for targeting as they had been found to be especially amenable to transactivation when targeted with dCas9 based synthetic transcription factors (Perez-Pinera et al., 2013). Six crRNAs were designed to target each of the three associated promoters. The crRNAs were designed to utilise a ‘TTV’ PAM sequence within a window 50 to 300 nt upstream of the transcription start site (TSS), identified using Fantom5 (Lizio et al., 2015) (Figure 3A). This window was selected as previous work had shown maximal transactivation of endogenous genes was obtained when targeting this window with a Cas9 derived synthetic transcription factor (Gilbert et al., 2014).

**Figure 3.**
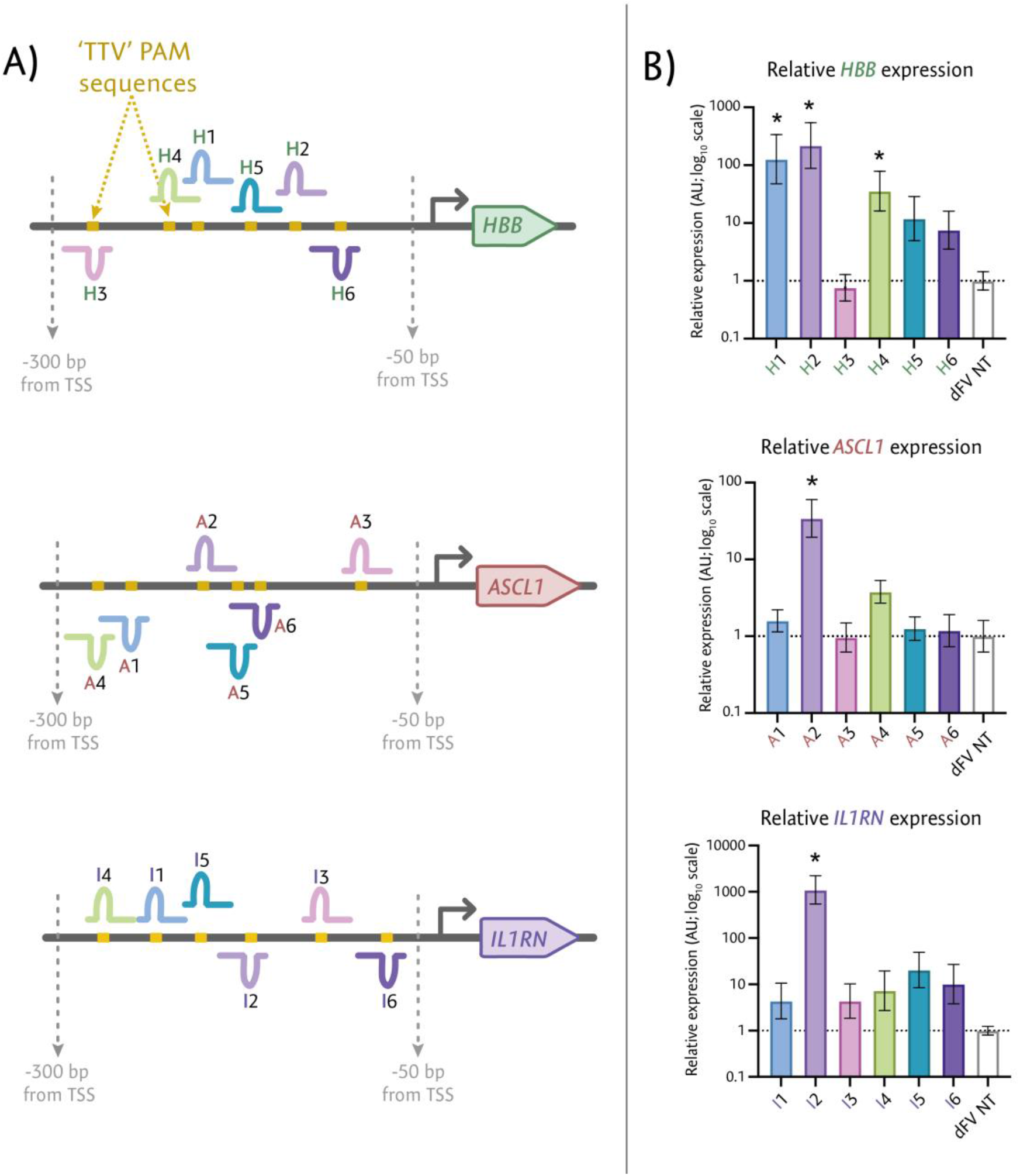
Testing single crRNAs for endogenous gene activation. **A)** dFnCas12a-VPR was screened for activity targeting 3 endogenous genes; *HBB, ASCL1* and *IL1RN* in HEK293 cells using single crRNAs. The promoter of each gene was targeted with six single crRNAs and transcriptional upregulation was compared to a non-targeting (NT) crRNA. **B)** The change in transcript abundance was then measured using qRT-PCR, with results shown for the three biological replicates. Stars (*) show results with a P value < 0.05 after one-way ANOVA followed by a post-hoc Dunnetts test to compare the expression of each targeting crRNA relative to the non-targeting negative control.

The crRNA plasmids were individually transfected alongside dFnCas12a-VPR into HEK293 cells before extracting the total RNA three days post transfection followed by qRT-PCR. When assessing the gene expression across all three genes, at least one crRNA for each promoter showed significant transactivation (Figure 3B). Statistically significant transactivation was observed for; *HBB* crRNA 1 (P = 0.0028), *HBB* crRNA 2 (P = 0.0011), *HBB* crRNA 4 (P = 0.0253), *ASCL1* crRNA 2 (P = 0.0003) and *IL1RN* crRNA 2 (P = 0.0002).

### 4 – Targeting multiple crRNAs enhances transactivation with evidence for synergy

Having shown that targeting dFnCas12a-VPR using single crRNAs was sufficient for transactivation, we next explored if the targeting of multiple crRNAs to the same promoter further enhanced gene expression synergistically. For each gene, the two individual crRNAs that showed the highest fold up-regulation were screened for transactivation, comparing their activity when co-transfected compared to individually transfected. For *HBB* and *IL1RN* a further crRNA pair was selected from crRNAs which had shown either weak or no significant transactivation (H4 + H5 and I4 + I6). To enable assessment of the relative impact of delivering two crRNAs compared to individual crRNAs, equimolar concentrations of crRNA plasmids were delivered to HEK293 cells.

Analysis by qRT-PCR showed the mean increase in mRNA abundance for the co-transfected condition was consistently higher than the most active individual crRNA (Figure 4A). When a two tailed t-test between the co-transfected and most active individual crRNA conditions was performed, we saw a significant increase in mRNA abundance for; *HBB* crRNA 1 + 2 (P = 0.017), *ASCL1* crRNA 2 + 4 (P = 0.001), and *IL1RN* crRNA 4 + 6 (P = 0.005). We also observed a non-significant increase in mRNA abundance for *HBB* crRNA 4 + 5 (P = 0.0875). For one of the tested crRNA pairs *(IL1RN* crRNA 2 and 5) the spacer sequences had partial complementarity (12 nucleotides). This may explain the small decrease in mRNA abundance observed when comparing co-transfection to *IL1RN* crRNA 2 individually (non-significant, P = 0.117). As a result, the *IL1RN* crRNA 2 + 5 pair was excluded from subsequent analysis.

**Figure 4.**
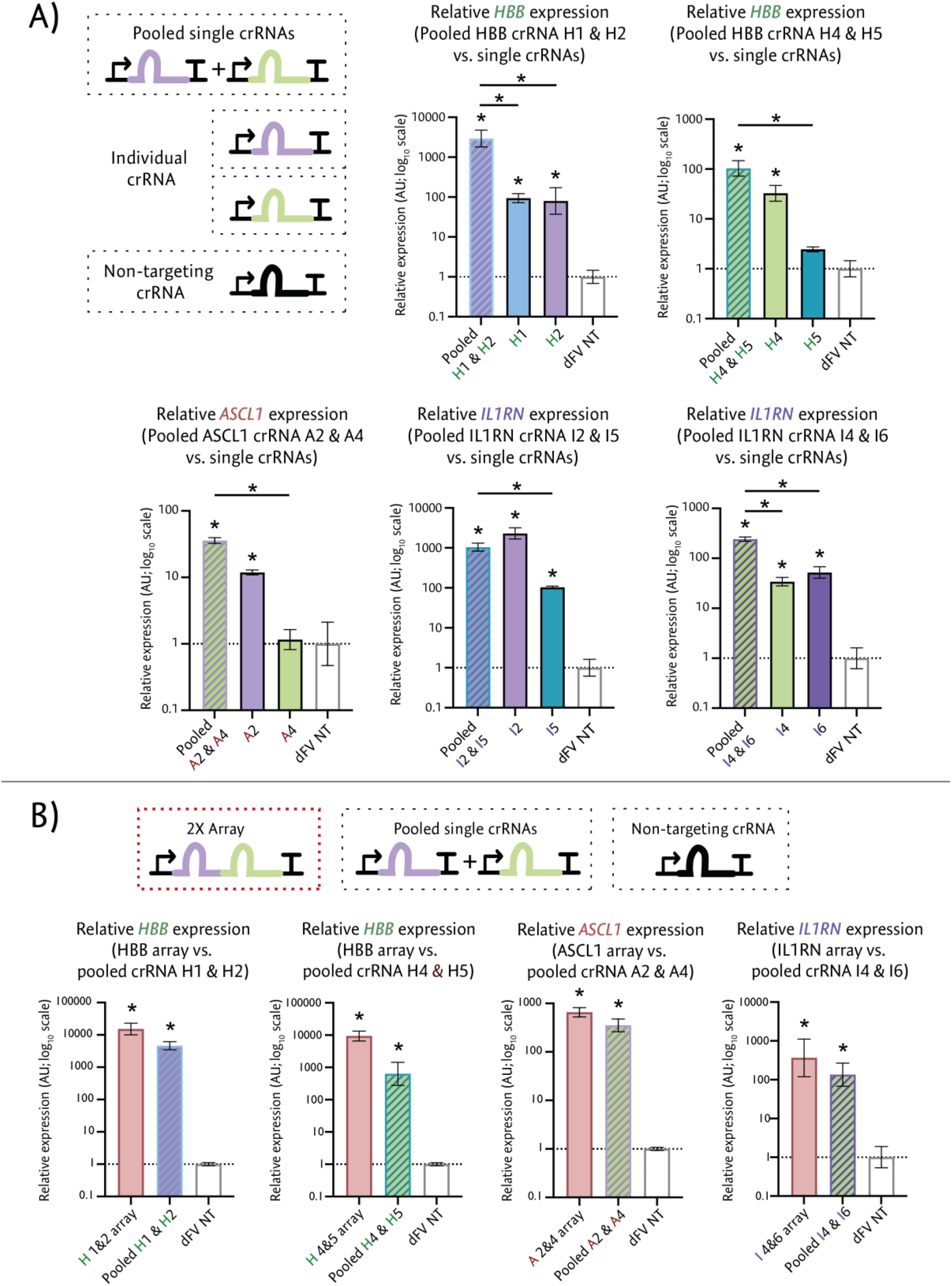
Enhanced activation observed with co-expression of targeting crRNAs. **A)** The most active individual crRNAs (from Figure 3) were delivered individually or co-transfected into HEK293 cells. The RNA abundance for each targeted gene was then measured by qRT-PCR and normalised to RNA expression when dFnCas12a-VPR was delivered with a non-targeting (NT) crRNA. The pair *IL1RN* crRNA 2 and *IL1RN* crRNA 5 were excluded as the spacer sequences overlapped. **B)** The crRNA pairs were incorporated into a single 2-crRNA array and the activity of this array was screened compared to co-transfection of the single crRNAs. Results from three biological replicates were measured by qRT-PCR and normalised to delivery with a non-targeting crRNA, with stars (*) showing results with a P value < 0.05.

As one of the advantages of Cas12a is the capacity to process crRNA arrays, we sought to test whether 2-crRNA arrays, consisting of a pair of crRNAs in tandem, could be utilised by dFnCas12a-VPR for transactivating target genes and whether these short arrays would enable increased or synergistic activation compared to the delivery of individual crRNAs. To test this, 2-crRNA arrays were constructed using the most active crRNA pairs from the preceding experiments. Three days after transfection into HEK293 cells, the fold upregulation induced using these arrays was tested compared to the co-transfected crRNAs and a non-targeting crRNA control. We consistently observed that the crRNA arrays performed as well if not better than the co-transfected crRNAs, with a higher mean fold upregulation for the arrays compared with the co-transfected crRNAs in all cases (Figure 4B).

We subsequently sought to test the crRNA arrays for evidence of synergistic transactivation. Synergy is here defined as showing greater transactivation than would be expected from adding the transactivation caused by each individual crRNA. To test this a hypothetical additive distribution was calculated by adding the distributions for the single crRNA conditions (Supplementary Note 1). Two tailed t-tests were then used to check whether significantly greater activation was seen for the array conditions than their respective hypothetical additive distributions. We saw that synergy was indeed observed for two of the cases, *HBB* (1+2) array (P = 0.0027) and *HBB* (4+5) array (P = 0.0027) (Supplementary Figure 1). Significance was not achieved for the *ASCL1* array (P = 0.1891) or the IL1RN array (P = 0.1069).

### 5 – Multiplexed activation from crRNA arrays

Having observed that transactivation of individual genes could be achieved using arrays with dFnCas12a-VPR, we next sought to test longer arrays designed to target multiple genes simultaneously, while exploring the impact of crRNA order within the array on activity. To achieve this, the most active crRNA arrays from the previous experiment (Figure 5A) were utilised for the generation of 6-crRNA arrays. Six different arrays were designed with three pairs of crRNAs, such that each pair targeted one of three promoters (Figure 5A). Using this design, all six combinations could be explored to not only test for significant multiplexed activation but also test whether changing the position or flanking crRNA sequences impacted the capacity of a crRNA array to induce transactivation. When screening mRNA abundance for each of the three genes, the results showed that each crRNA array was able to significantly up-regulate transcription for every gene (Figure 5B).

**Figure 5.**
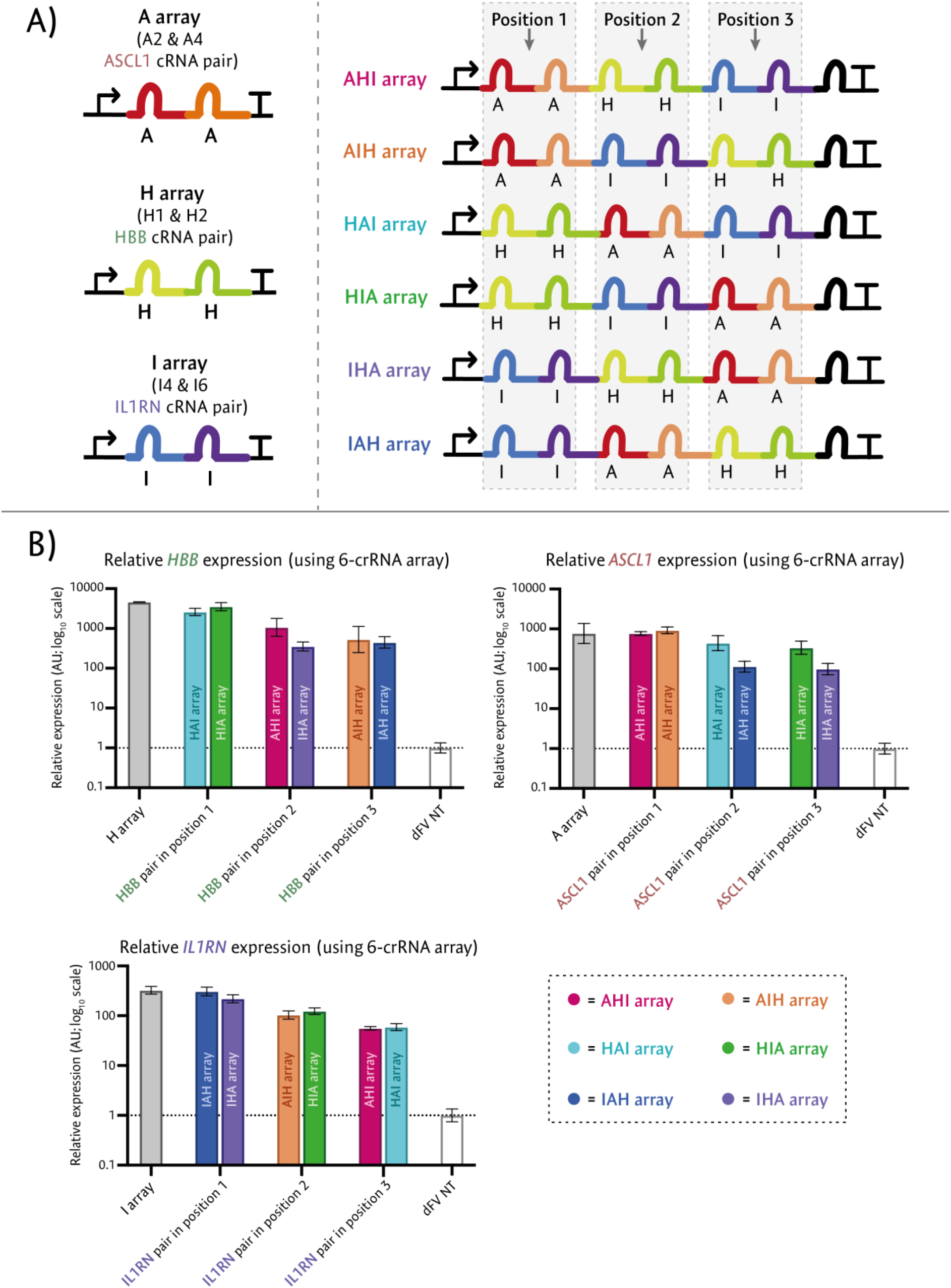
Multiplexed activation of endogenous genes. **A)** The left panel shows a schematic of the most active crRNA array for targeting each of the three genes from Figure 4B. The right panel shows the designs of the combinatorial 6-crRNA arrays for screening for multiplexed activation. The six arrays were designed to ensure each of the pairs of crRNAs would be present in the first and second, third and fourth or fifth and sixth positions within the 6-crRNA arrays. **B)** The six arrays were separately transfected into HEK293 cells alongside dFnCas12a-VPR and the expression for each of the three targeted genes was assessed by qRT-PCR, normalising to the expression with a non-targeting crRNA. The results from the three biological replicates are displayed based upon the position of the respective targeting crRNAs within the 6-crRNA arrays.

### 6 – Modest order dependent array activity observed for individual and paired crRNA

For all three genes targeted, there appeared to be a clear correlation between the position of the targeting crRNAs within the arrays and the fold up-regulation. More specifically when the gene targeting crRNAs were positioned closer to the 3’ end of the array, increases in mRNA abundance were consistently diminished across all three genes tested (Figure 5B). When simple linear regression was performed, a weak inverse correlation between the targeting crRNA position within an array and gene activation of the targeted gene was observed for *ASCL1* and *HBB* (R^2^ = 0.4160, P = 0.0039 and R^2^ = 0.4876, P = 0.0013 respectively) and a strong inverse correlation was observed for *IL1RN* (R^2^ = 0.8457, P < 0.0001) (Supplementary Figure 1).

To further investigate this phenomenon, a series of crRNA arrays were generated where each array possessed a single targeting crRNA and five non-targeting crRNAs. The non-targeting crRNAs were rationally designed from different randomly generated 20 nucleotide sequences, that showed no perfect matches against the human genome.

Six different versions of the array were generated so that all positions of the targeting crRNA (*ASCL1* crRNA 2) within the array could be tested (Figure 6A). The capacity of the each crRNA array to up-regulate *ASCL1* expression was then measured as previously described. The results showed a reduction in transactivation of *ASCL1* as the targeting crRNA was positioned closer to the 3’ of the array, with the highest mean fold-upregulation (103-fold) when the crRNA was at the first (most 5’) position and the lowest fold up-regulation (29fold) when the crRNA was at the last position (most 3’) (Figure 6B). When simple linear regression was performed, a weak reduction in activity when the targeting crRNAs were positioned towards the 3’ of the arrays was observed (R^2^ = 0.3671, P = 0.0077) (Supplementary Figure 2).

**Figure 6.**
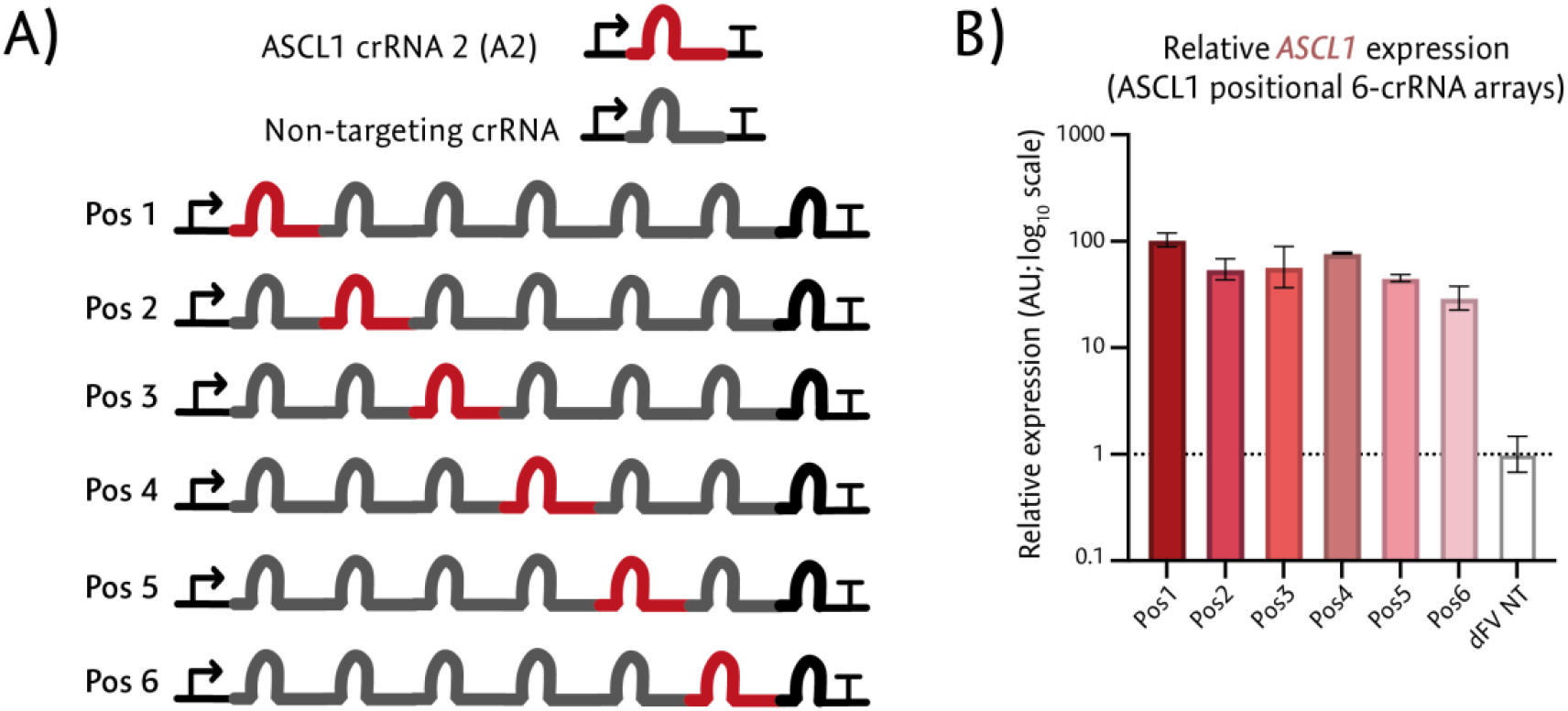
Position dependent activity within pol3 derived crRNA arrays. **a)** Design of arrays constructed for testing the impact of position for a single targeting crRNA within a 6-crRNA array. **b)** qRT-PCR analysis of *ASCL1* mRNA abundance for the six arrays after transfection into HEK293 cells, with the graph showing results from the three biological replicates.

## Discussion

Here we have shown the first application of engineering FnCas12a derived synthetic transcription factors in mammalian cells. The key advantage of the Fn variant is the simpler PAM sequence ‘KYTV’ when compared to the commonly utilised As or Lb variants ‘TTTV’. This translates to being able to target on average every 21 nt as opposed to on average every 85 nt, highly comparable to the Cas9 PAM sequence ‘NGG’ which enables targeting on average every 16 nt. This allows much denser targeting, with more potential targets within any given length of DNA. This is of particular interest for transactivation of target genes as we have also highlighted that delivery of multiple active crRNAs targeting the same promoter region further enhances up-regulation.

We found that dFnCas12a-VPR can be used for multiplexed transactivation of three different genes from a single transcript. We have consistently observed a reduction in activity for crRNAs positioned closer to the 3’ of crRNA arrays expressed from the U6 pol 3 promoter. This information can inform design constraints when targeting multiple genes for upregulation from a single array. In particular, arrays can be designed to express crRNAs closer to the 5’ end of an array where they target genes that are challenging to upregulate or where higher overexpression is desired. Conversely, crRNAs targeting genes that are easier to up-regulate or where lower over-expression is desired can be positioned closer to the 3’ end of an array.

One likely explanation for the reduction of activity as the crRNA is positioned closer to the 3’ end of the array is that crRNA abundance is reduced. Zetsche *et al*. showed with RNA-seq data a decreased crRNA abundance towards the 3’ of a 3-crRNA array when expressed in HEK293 cells (Zetsche et al., 2017). This may be due to the presence of a weak non-canonical Pol III terminator sequence ‘TTTCT’ (Orioli et al., 2011) within all of the direct repeats within the crRNA array.

Through this expansion of the Cas12a toolkit researchers should be able to more easily simultaneously transactivate multiple genes, with the capacity to densely target multiple promoters. In addition, the compact nature of the crRNA array both facilitates cheap and easy assembly using short oligo-based assembly strategies. Furthermore the compact arrays expand the potential of CRISPR systems when considering AAV viral delivery for therapeutic applications (which has a packaging size limit of 4.2Kbp) when paired with split Cas12a strategies (Kempton et al., 2020).

Whilst previous work has explored the role of PAM selection (Jacobsen et al., 2020) and spacer sequence choice (Creutzburg et al., 2020) on crRNA activity, to our knowledge this is the first time where order dependent activity for hU6 expressed crRNA arrays have been shown. This is of key relevance to researchers as U6 promoters naturally highly express short non-coding RNA, with a defined termination sequence of five thymidines. As such the majority of crRNA expression plasmids utilise this promoter and the findings of order dependent activity will have relevance to researchers working with Cas12a or derived synthetic transcription factors. In particular the order dependent activity of crRNAs within an array also opens up the possibility of diversifying the fold change of transactivation of different genes within genetic networks and pathways.

This may open up opportunities for streamlined manipulation of pathways where for example the promoters of multiple genes within a pathway can be targeted by the same crRNA but in different orderings. Subsequent sequencing of high production strains can reveal the enrichment of order for specific crRNAs, which in turn reveals when higher transactivation or reduced transactivation of a specific gene with a pathway are being selected for. This in turn can help to reveal key bottlenecks or toxicities that emerge within a pathway. A similar approach can be considered for processes such as cell reprogramming and indeed any process where transcriptional modulation of a gene network can be correlated with a phenotype.

## Materials and Methods

### Plasmid construction

Isothermal mutagenesis was used to introduce amino acid substitutions; D908A for AsCas12a, D917A for FnCas12a and D832A for LbCas12a. Mutation of this highly conserved amino acid has previously been shown to abolish DNase activity (Zetsche et al., 2015). The VPR transactivation domain was subcloned onto the 3’ of each dCas12a from dCas9-VPR, restriction ligation.

Single crRNA plasmids and crRNA arrays were generated by first annealing oligos ordered from IDT (Integrated DNA Technologies). The annealed oligos were then ligating into the BpiI (Thermo Scientific cat #ER1012) digested pU6 plasmid backbone using 1μl of T4 PNK (NEB cat #M0201L) and 1μl of T4 ligase (NEB cat #M0202L) in a 20μl reaction, incubated at 37°C for 30 minutes before transforming into *E.coli*.

### Cell culturing and transfection

HEK293 cells were cultured in DMEM (Gibco; Life Technologies) with 10% FBS (Gibco; Life Technologies), 4mM glutamine and 1% penicillin-streptomycin (Gibco; Life Technologies). Transfections were carried out a day after seeding ~200,000 cells per well into 24 well plates. Transfections were performed using either lipofectamine 2000 (luciferase assays) or GenJet In vitro transfection reagent (qRT-PCRs). For the initial luciferase assays 200ng of the synthetic transcription factor was transfected with 100ng of the gRNA/crRNA plasmids. For the qRT-PCR assays 500ng of the synthetic transcription factor was transfected with 250ng of the gRNA/crRNA plasmids.

### Dual luciferase assay

HEK293 cells were transfected with a Firefly luciferase reporter construct, Renilla luciferase normalising construct and the respective synthetic transcription factor construct, with or without a targeting crRNA/gRNA plasmid. Firefly luciferase expression and Renilla luciferase expression were then assessed using the dual-luciferase kit (Promega E1910) with measurements carried out with the Modulus II microplate reader (Turner Biosystems). In all cases cells were lysed in passive lysis buffer 48 hours after transfection.

### RNA extraction and cDNA generation

72 hours after transfection cells were harvested and RNA extraction was performed using E.Z.N.A Total RNA Kit 1 (Omega Biotek cat #R6834-01). cDNA generation was performed using SuperScript IV Reverse Transcriptase (Invitrogen). 1 μg of RNA was mixed with 1μl of 50μM oligo d(T)20 (IDT), 1μl of 10mM dNTP mix (Promega cat #U1240) and DEPC water up to a final volume of 13μl in a PCR tube and incubated at 65°C for 5 minutes then on ice for 1 minute. The following components were added to each sample: 4μl of SuperScript IV Reverse Transcriptase buffer, 1μl of 0.1 M DTT, 1ul of dH2O, 0.5μl of RiboLock RNase Inhibitor (Invitrogen) and 0.5μl of SuperScript IV Reverse Transcriptase (Invitrogen). The reactions were then incubated at 52°C for 10 minutes followed by 80°C for 10 minutes and holding at 4°C.

Unless otherwise stated, cells were harvested three days post transfection using E.Z.N.A Total RNA Kit 1 (Omega Biotek cat #R6834-01) according to the manufacturer’s instructions. The concentration and RNA quality was assessed using the nanodrop.

1μl of 50μM oligo d(T)20 (IDT) and 1μl of 10mM dNTP mix (Promega cat #U1240) were combined with 1 μg of RNA before adding dd H20 up to a final volume of 13μl. cDNA was then generated using SuperScript IV Reverse Transcriptase (Thermo Fisher cat #18090050) according to the manufacturer’s instructions using 0.5μl of RiboLock RNase Inhibitor (Thermo Scientific cat #EO0381) and 0.5μl of SuperScript IV Reverse Transcriptase per sample.

### qRT-PCR

The 96 well qPCR reaction plates were set up using the Power SYBR Green qPCR mix (Thermo Fisher cat #4367659) according to the manufacturer’s protocol, using a 10μl total reaction volume. The 96 well qPCR plate was run on the StepOnePlus real-time PCR machine (Thermo Fisher cat #4376600). Cycle threshold (Ct) values were calculated on the StepOnePlus PCR machine software and further analysed using the statistical analysis software, Prism 8.

384 well qPCR reaction plates for the multiplexed activation of endogenous genes (Figure 5 and 6) were set up using the Brilliant II SYBR master mix (Agilent cat #600828) according to the manufacturer’s protocol, using a 4 ul total reaction volume. Samples were loaded on a 384 multiwell plate (Roche cat #04729749001) and ran on the Lightcycler 480 qPCR machine. The Ct values were calculated on the Lightcycler software and further analysed using the statistical analysis software, Prism 8.

## Acknowledgements

This work was supported by the BBSRC (BB/M018040/1). We wish to thank Dr Tessa Moses for her discussion and advice, Sam Haynes for advice on statistical approaches as well as Dr Mathew Dale, Jessica Birt and Trevor Ho for their comments on the initial manuscript.

## Notes

### Competing Interest Statement

The authors have declared no competing interest.

## References

Anders, C., Niewoehner, O., Duerst, A., Jinek, M., 2014. Structural basis of PAM-dependent target DNA recognition by the Cas9 endonuclease. Nature 513, 569–573. https://doi.org/10.1038/nature13579

Becskei, A., 2020. Tuning up Transcription Factors for Therapy. Molecules 25. https://doi.org/10.3390/molecules25081902

Campa, C.C., Weisbach, N.R., Santinha, A.J., Incarnato, D., Platt, R.J., 2019. Multiplexed genome engineering by Cas12a and CRISPR arrays encoded on single transcripts. Nat Methods 16, 887–893. https://doi.org/10.1038/s41592-019-0508-6

Chakraborty, S., Ji, H., Kabadi, A.M., Gersbach, C.A., Christoforou, N., Leong, K.W., 2014. A CRISPR/Cas9-Based System for Reprogramming Cell Lineage Specification. Stem Cell Reports 3, 940–947. https://doi.org/10.1016/j.stemcr.2014.09.013

Creutzburg, S.C.A., Wu, W.Y., Mohanraju, P., Swartjes, T., Alkan, F., Gorodkin, J., Staals, R.H.J., van der Oost, J., 2020. Good guide, bad guide: spacer sequence-dependent cleavage efficiency of Cas12a. Nucleic Acids Res 48, 3228–3243. https://doi.org/10.1093/nar/gkz1240

Fonfara, I., Richter, H., Bratovič, M., Le Rhun, A., Charpentier, E., 2016. The CRISPR-associated DNA-cleaving enzyme Cpf1 also processes precursor CRISPR RNA. Nature 532, 517–521. https://doi.org/10.1038/nature17945

Gilbert, L.A., Horlbeck, M.A., Adamson, B., Villalta, J.E., Chen, Y., Whitehead, E.H., Guimaraes, C., Panning, B., Ploegh, H.L., Bassik, M.C., Qi, L.S., Kampmann, M., Weissman, J.S., 2014. Genome-Scale CRISPR-Mediated Control of Gene Repression and Activation. Cell 159, 647–661. https://doi.org/10.1016/j.cell.2014.09.029

Gilbert, L.A., Larson, M.H., Morsut, L., Liu, Z., Brar, G.A., Torres, S.E., Stern-Ginossar, N., Brandman, O., Whitehead, E.H., Doudna, J.A., Lim, W.A., Weissman, J.S., Qi, L.S., 2013. CRISPR-Mediated Modular RNA-Guided Regulation of Transcription in Eukaryotes. Cell 154, 442–451. https://doi.org/10.1016/j.cell.2013.06.044

Hilton, I.B., D’Ippolito, A.M., Vockley, C.M., Thakore, P.I., Crawford, G.E., Reddy, T.E., Gersbach, C.A., 2015. Epigenome editing by a CRISPR-Cas9-based acetyltransferase activates genes from promoters and enhancers. Nature Biotechnology 33, 510–517. https://doi.org/10.1038/nbt.3199

Jacobsen, T., Ttofali, F., Liao, C., Manchalu, S., Gray, B.N., Beisel, C.L., 2020. Characterization of Cas12a nucleases reveals diverse PAM profiles between closely-related orthologs. Nucleic Acids Res 48, 5624–5638. https://doi.org/10.1093/nar/gkaa272

Jinek, M., Chylinski, K., Fonfara, I., Hauer, M., Doudna, J.A., Charpentier, E., 2012. A Programmable Dual-RNA–Guided DNA Endonuclease in Adaptive Bacterial Immunity. Science 337, 816–821. https://doi.org/10.1126/science.1225829

Kempton, H.R., Goudy, L.E., Love, K.S., Qi, L.S., 2020. Multiple Input Sensing and Signal Integration Using a Split Cas12a System. Molecular Cell 78, 184–191.e3. https://doi.org/10.1016/j.molcel.2020.01.016

Kim, D., Kim, J., Hur, J.K., Been, K.W., Yoon, S., Kim, J.-S., 2016. Genome-wide analysis reveals specificities of Cpf1 endonucleases in human cells. Nature Biotechnology 34, 863–868. https://doi.org/10.1038/nbt.3609

Kim, H.K., Song, M., Lee, J., Menon, A.V., Jung, S., Kang, Y.-M., Choi, J.W., Woo, E., Koh, H.C., Nam, J.-W., Kim, H., 2017. *In vivo* high-throughput profiling of CRISPR– Cpf1 activity. Nature Methods 14, 153–159. https://doi.org/10.1038/nmeth.4104

Kleinjan, D.A., Wardrope, C., Sou, S.N., Rosser, S.J., 2017. Drug-tunable multidimensional synthetic gene control using inducible degron-tagged dCas9 effectors. Nat Commun 8, 1–9. https://doi.org/10.1038/s41467-017-01222-y

Krawczyk, K., Scheller, L., Kim, H., Fussenegger, M., 2020. Rewiring of endogenous signaling pathways to genomic targets for therapeutic cell reprogramming. Nat Commun 11, 608. https://doi.org/10.1038/s41467-020-14397-8

Lizio, M., Harshbarger, J., Shimoji, H., Severin, J., Kasukawa, T., Sahin, S., Abugessaisa, I., Fukuda, S., Hori, F., Ishikawa-Kato, S., Mungall, C.J., Arner, E., Baillie, J.K., Bertin, N., Bono, H., de Hoon, M., Diehl, A.D., Dimont, E., Freeman, T.C., Fujieda, K., Hide, W., Kaliyaperumal, R., Katayama, T., Lassmann, T., Meehan, T.F., Nishikata, K., Ono, H., Rehli, M., Sandelin, A., Schultes, E.A., ’t Hoen, P.A.C., Tatum, Z., Thompson, M., Toyoda, T., Wright, D.W., Daub, C.O., Itoh, M., Carninci, P., Hayashizaki, Y., Forrest, A.R.R., Kawaji, H., FANTOM consortium, 2015. Gateways to the FANTOM5 promoter level mammalian expression atlas. Genome Biol. 16, 22. https://doi.org/10.1186/s13059-014-0560-6

Maeder, M.L., Linder, S.J., Cascio, V.M., Fu, Y., Ho, Q.H., Joung, J.K., 2013. CRISPR RNA–guided activation of endogenous human genes. Nature Methods 10, 977–979. https://doi.org/10.1038/nmeth.2598

Mali, P., Yang, L., Esvelt, K.M., Aach, J., Guell, M., DiCarlo, J.E., Norville, J.E., Church, G.M., 2013. RNA-Guided Human Genome Engineering via Cas9. Science 339, 823–826. https://doi.org/10.1126/science.1232033

Nakamura, M., Srinivasan, P., Chavez, M., Carter, M.A., Dominguez, A.A., La Russa, M., Lau, M.B., Abbott, T.R., Xu, X., Zhao, D., Gao, Y., Kipniss, N.H., Smolke, C.D., Bondy-Denomy, J., Qi, L.S., 2019. Anti-CRISPR-mediated control of gene editing and synthetic circuits in eukaryotic cells. Nat Commun 10. https://doi.org/10.1038/s41467-018-08158-x

Orioli, A., Pascali, C., Quartararo, J., Diebel, K.W., Praz, V., Romascano, D., Percudani, R., van Dyk, L.F., Hernandez, N., Teichmann, M., Dieci, G., 2011. Widespread occurrence of non-canonical transcription termination by human RNA polymerase III. Nucleic Acids Res 39, 5499–5512. https://doi.org/10.1093/nar/gkr074

Pandelakis, M., Delgado, E., Ebrahimkhani, M.R., 2020. CRISPR-Based Synthetic Transcription Factors In Vivo: The Future of Therapeutic Cellular Programming. Cell Systems 10, 1–14. https://doi.org/10.1016/j.cels.2019.10.003

Perez-Pinera, P., Kocak, D.D., Vockley, C.M., Adler, A.F., Kabadi, A.M., Polstein, L.R., Thakore, P.I., Glass, K.A., Ousterout, D.G., Leong, K.W., Guilak, F., Crawford, G.E., Reddy, T.E., Gersbach, C.A., 2013. RNA-guided gene activation by CRISPR-Cas9-based transcription factors. Nat. Methods 10, 973–976. https://doi.org/10.1038/nmeth.2600

Tak, Y.E., Kleinstiver, B.P., Nuñez, J.K., Hsu, J.Y., Horng, J.E., Gong, J., Weissman, J.S., Joung, J.K., 2017. Inducible and multiplex gene regulation using CRISPR–Cpf1-based transcription factors. Nature Methods 14, 1163–1166. https://doi.org/10.1038/nmeth.4483

Tu, M., Lin, L., Cheng, Y., He, X., Sun, H., Xie, H., Fu, J., Liu, C., Li, J., Chen, D., Xi, H., Xue, D., Liu, Q., Zhao, J., Gao, C., Song, Z., Qu, J., Gu, F., 2017. A ‘new lease of life’: FnCpf1 possesses DNA cleavage activity for genome editing in human cells. Nucleic Acids Res 45, 11295–11304. https://doi.org/10.1093/nar/gkx783

Zetsche, B., Gootenberg, J.S., Abudayyeh, O.O., Slaymaker, I.M., Makarova, K.S., Essletzbichler, P., Volz, S.E., Joung, J., van der Oost, J., Regev, A., Koonin, E.V., Zhang, F., 2015. Cpf1 is a single RNA-guided endonuclease of a class 2 CRISPR-Cas system. Cell 163, 759–771. https://doi.org/10.1016/j.cell.2015.09.038

Zetsche, B., Heidenreich, M., Mohanraju, P., Fedorova, I., Kneppers, J., DeGennaro, E.M., Winblad, N., Choudhury, S.R., Abudayyeh, O.O., Gootenberg, J.S., Wu, W.Y., Scott, D.A., Severinov, K., van der Oost, J., Zhang, F., 2017. Multiplex gene editing by CRISPR-Cpf1 through autonomous processing of a single crRNA array. Nat Biotechnol 35, 31–34. https://doi.org/10.1038/nbt.3737

