## Supplementary Figures for "Multiplexed activation in mammalian cells using dFnCas12a-VPR"

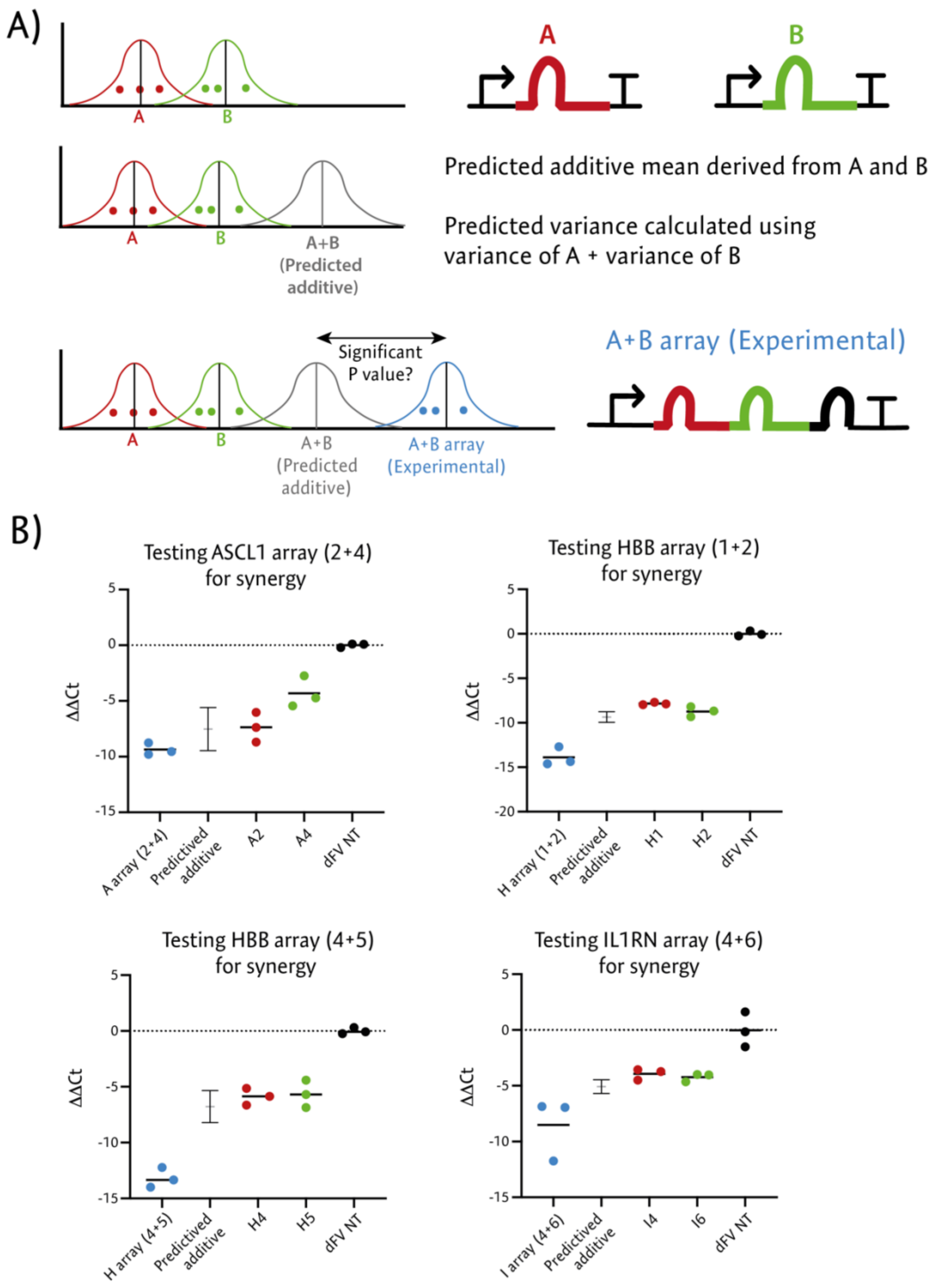


**Supplementary Figure 1 – Testing crRNA arrays for synergistic transactivation**
The ΔΔCT values used to calculate the relative mRNA expression shown in figure 5B were used to calculate a hypothetical additive distribution using the approach outlined in Supplementary Note 1. The error bars denote the predicted standard deviation and the grey bar in the centre denotes the predicted mean. A one tailed t-test was then performed between each array and the associated hypothetical additive condition to test for synergistic (greater than additive) transactivation.

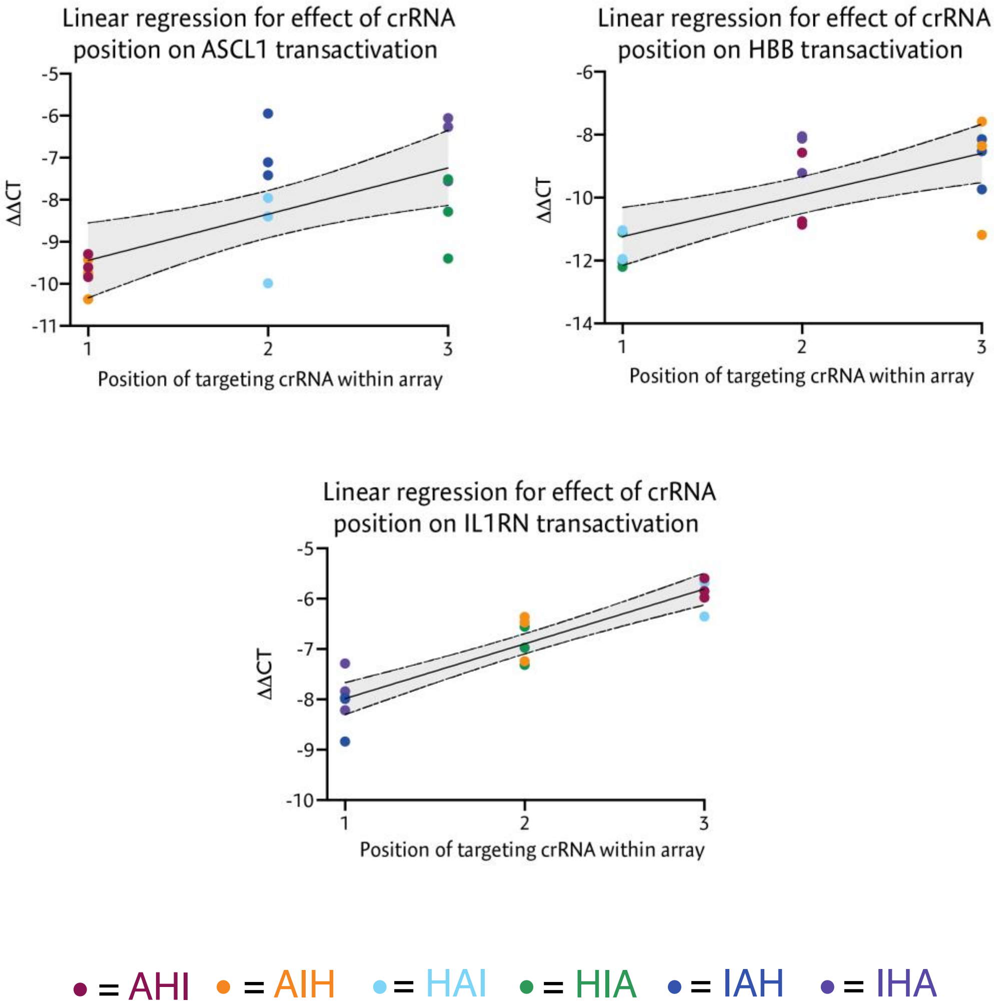


**Supplementary Figure 2 – Simple linear regression for multiplexing crRNA array**
The ΔΔCT values used to calculate the relative mRNA expression shown in figure 6 have been tested for a linear trend using linear regression analysis. For each of the three targeted genes, a significant increase in ΔΔCT (representing a decrease in relative mRNA abundance) is observed as you move the targeting crRNA pair from position 1 (most 5') towards position 3 (most 3') within the crRNA array ($R^2$ = 0.4160, P = 0.0039 for *ASCL1*; $R^2$ = 0.4876, P = 0.0013 for HBB and $R^2$ = 0.8457, P < 0.0001 for IL1RN.

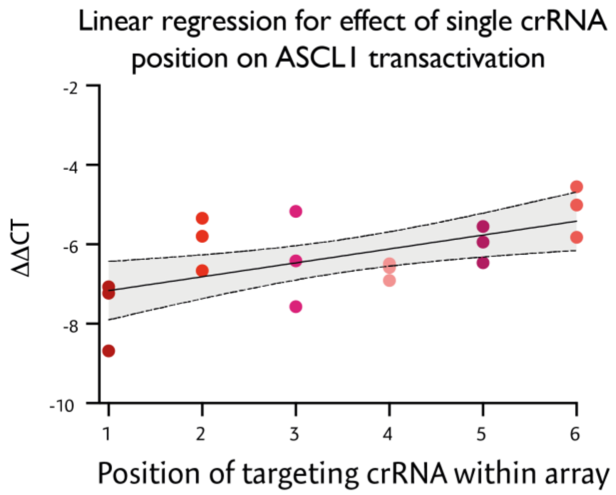


**Supplementary Figure 3 – Simple linear regression for single active crRNA within array**
The ΔΔCT values used to calculate the relative mRNA expression shown in figure 7 have been tested for a linear trend using linear regression analysis. A significant increase in ΔΔCT (representing a decrease in relative mRNA abundance) is observed as you move the targeting crRNA from position 1 (most 5') to position 6 (most 3') ($R^2$ = 0.3671, P = 0.0077).

### Supplementary notes

#### Supplementary Note 1

The means and standard deviations for the predicted additive ΔΔCT distributions shown in supplementary figure 1B were derived by inferring the expected additive distribution for the fold change of mRNA when both targeting crRNA were present. The inferred mean ΔΔCT values was calculated by adding the geometric means for each of the two individual crRNA mRNA fold changes. The inferred variance was calculated by adding the variances for the CT values for each of the individual crRNA. From this variance the standard deviation was calculated by taking the square root of this value.
